# ZBTB38 is dispensable for hematopoiesis and antibody responses

**DOI:** 10.1101/2020.06.11.145839

**Authors:** Rachel Wong, Deepta Bhattacharya

## Abstract

Members of the broad complex, tram track, bric-a-brac and zinc finger (BTB-ZF) family of transcription factors, such as BCL-6, ZBTB20, and ZBTB32, regulate antigen-specific B cell differentiation, plasma cell longevity, and the duration of antibody production. We found that ZBTB38, a different member of the BTB-ZF family that binds methylated DNA at CpG motifs, is highly expressed by germinal center B cells and plasma cells. To define the functional role of ZBTB38 in B cell responses, we generated mice conditionally deficient in this transcription factor. Germinal center B cells lacking ZBTB38 dysregulated very few genes relative to wild-type and heterozygous littermate controls. Accordingly, mice with hematopoietic-specific deletion of *Zbtb38* showed normal germinal center B cell numbers and antibody responses following immunization with hapten-protein conjugates. Memory B cells from these animals functioned normally in secondary recall responses. Despite expression of ZBTB38 in hematopoietic stem cells, progenitors and mature myeloid and lymphoid lineages were also present in normal numbers in mutant mice. These data demonstrate that ZBTB38 is dispensable for hematopoiesis and antibody responses. These conditional knockout mice may instead be useful in defining the functional importance of ZBTB38 in other cell types and contexts.

## Introduction

Antibody responses following infections or vaccinations are initiated by a series of B cell activation steps and fate decisions [1]. Upon recognition of cognate antigens and other stimulatory signals, B cells grow in size, express a panel of activation markers, begin to proliferate, and a subset undergoes immunoglobulin isotype-switching [2]. In T cell-dependent responses, B cells then differentiate either into antibody-secreting plasma cells or into germinal center B cells. Germinal centers are the sites in which somatic hypermutation and affinity maturation occur and are under substantial replicative and DNA damage-induced stress. Germinal centers eventually produce long-lived plasma cells (LLPCs) and memory B cells (MBCs), which have distinct antigen specificities and mediate different aspects of immunity [3]. LLPCs constitutively secrete antibodies and are important for providing protection against re-infection by the same pathogen. Memory B cells, on the other hand, can only provide protection after re- activation by a cognate antigen through rapid differentiation into plasma cells. Recent studies have identified the broad complex, tram track, bric-a-brac and zinc finger (BTB-ZF) family of transcription factors as key regulators in B cell development. BTB-ZF family members bind DNA through its C-terminal zinc finger domains and recruit SMRT co-repressors and histone deacetylases to N-terminal BTB/POZ domains [4-8]. Family members that regulate distinct aspects of B cell-mediated immunity include BCL-6, ZBTB20, and ZBTB32. BCL-6 is important for germinal center (GC) formation [9], ZBTB20 promotes plasma cell lifespan and durable immunity in an adjuvant dependent manner [10, 11], and ZBTB32 restricts memory B cell recall responses [12, 13]. Other as-yet uncharacterized BTB-POZ members may regulate different aspects of B cell responses [4].

ZBTB38, also known as CIBZ (CtBP-interacting BTB zinc finger protein), is another member of the BTB-ZF family and can function either as a transcriptional repressor or activator [14]. ZBTB38-mediated transcriptional regulation occurs by binding primarily to specific methylated CpG sequences and recruiting repressors or activators [15-19]. ZBTB38 has been shown to repress overall transcription by inhibiting expression of MCM10, a component of the pre-replication complex [20]. Additionally, ZBTB38 can, directly or indirectly, regulate cell cycle progression, cellular differentiation, and apoptosis [21-25]. Here, we demonstrate that, despite high levels of expression in germinal center B lymphocytes and plasma cells, ZBTB38 deficiency does not impair primary or secondary antibody responses to T cell-dependent model immunogens.

## Materials and Methods

### Ethics statement

All procedures in this study were specifically approved and carried out in accordance with the guidelines set forth by the Institutional Animal Care and Use Committee at Washington University (approval 20140030) and at the University of Arizona (approval 17-266). Euthanasia was performed by administering carbon dioxide at 1.5L/minute into a 7L chamber until 1 minute after respiration ceased. After this point, cervical dislocation was performed to ensure death.

### Mice

All mice were housed and bred in pathogen-free facilities. C57BL6/N mice were obtained from the National Cancer Institute. B6.Cg-*Igh*^*a*^*Thy1*^*a*^*Gpi1*^*a*^ (*IgH*^*a*^) mice were obtained from Charles River Laboratories. *Zbtb38*^*fl/+*^ mice were generated by injecting C57Bl6/J pronuclear zygotes with ribonucleoparticles of Cas9 protein and guide RNAs targeting sites flanking exon 3 of *Zbtb38*, alongside oligonucleotide donors as homologous recombination donors spanning these same gRNA sites. Oligonucleotide substrates contained loxP sites to interrupt gRNA target sites to prevent Cas9 re-cutting after successful recombination. A single successful founder (out of 33 tested) was identified by PCR and then bred to C57Bl6/N mice for germline transmission. Mice have been maintained by backcrossing to C57Bl6/N animals. Animals will be made available at the Mutant Mouse Resource and Research Centers upon publication of this manuscript (B6N.B6J-*Zbtb38em1Dbhat*/Mmucd, Strain ID 66871). ZBTB38 f/f mice were crossed to CMV-Cre (Jackson Laboratory, stock no. 006054) or VavCre (Jackson Laboratory, stock no. 008610) and maintained as ZBTB38 f/f or ZBTB38 x CMV- or Vav-Cre where littermates were used as controls. The following primers were used to confirm recombination of the targeting plasmid: SP55.mZbtb38.5’genomic.F2 5’- CCAGGGATTCAGTCCTCAGCA-3’, SP55.mZbtb38.3’genomic.R2 5’-GCCTACCCCAAACCACACTAA-3’. The following primers were used for genotyping the *Zbtb38* allele: 5’LoxP forward 5’- TCTGAGTTCAAGGCCAGCTT-3’, 5’LoxP reverse 5’- TCTCCAAGCAGAAAGGGTGT-3’, and 3’LoxP reverse 5’- GGGTCGTTAGAGGATTCAGC-3’.

### Immunizations

Mice were immunized intraperitoneally with 100 μg NP-OVA (Biosearch), adjuvanted with Alhydrogel (Invivogen). NP-APC used for staining was made by conjugating allophycocyanin (Sigma-Aldrich) with 4-hydroxy-3- nitrophenylacetyl-O-succinimide ester (Biosearch Technologies).

### RNA extraction, cDNA synthesis, and qRT-PCR

Total RNA was extracted with TRIzol (Life technologies) and cDNA synthesized using Superscript III Reverse transcription kit with random hexamers (Life Technologies) according to manufacturer’s instructions. qRT-PCR was performed using SYBR Green PCR master mix (Applied Biosystems) on a Prism 7000 Sequence Detection System (Applied Biosystems). The primers used for *Zbtb38* are: forward 5’- AGAACCAAGGATTTCCGAGTG-3’ and reverse 5’-GATGGAGAGTACTGTGTCACTG-3’. *Zbtb38* transcript levels were normalized to 18S ribosomal RNA, forward 5’-CGGCTACCACATCCAAGGAA-3’ and reverse 5’-GCTGGAATTACCGCGGCT-3’ [26].

### RNA-sequencing

RNA from germinal center B cells was extracted with Macherey-Nagel Nucleospin XS kits. cDNA libraries were prepared by Novogene using SmartSeq v4 kits (Takara) and processed for paired-end PE150 RNA-sequencing on an Illumina Hiseq 4000 lane. For visualization of ZBTB38 transcripts, fastq files were mapped and aligned to the mm10 genome using HiSat2 and displayed using IgV [27, 28]. For quantification of gene expression differences, fastq files were mapped using vM17 annotation files from Gencode and transcript abundances were quantified by Salmon [29]. Differentially-expressed genes quantified by DESeq2 [30]. Volcano plots were displayed using Prism software (GraphPad). Data have been deposited to NCBI GEO and await an accession number.

### ELISA

ELISA plates (9018, Corning) were coated overnight at 4°C in 0.1 M sodium bicarbonate buffer, pH 9.5 containing 5 ug/mL of NP_16_- or NP_4_-BSA (Bioresearch Technologies). All other incubation steps were performed at room temperature for 1 hour. Wash steps were performed between each step using PBS + 0.05% Tween-20. Plates were blocked with PBS + 2% BSA followed by serial dilutions of serum. Serum was probed with 0.1 ug/mL of biotinylated anti-mouse IgG (715-065-151, Jackson ImmunoResearch Laboratories) and then detected with streptavidin conjugated horseradish peroxidase (554066, BD biosciences). Plates were developed using TMB (Dako, S1599) and neutralized with 2N H_2_SO_4_. Optical density (OD) values were measured at 450 nm. Serum endpoint titer was defined as the inverse dilution factor that is three standard deviations above background using one-phase decay measurements and Prism software (GraphPad Software).

### Adoptive transfer for recall responses

Splenocytes from NP-immunized *ZBTB38*^f/f^ or *ZBTB38*^f/f^ x VavCre mice were isolated and processed into single cell suspension. erythrocytes lysed using an ammonium chloride-potassium solution, and lymphocytes isolated by using a Hisopaque-1119 (Sigma-Aldrich) density gradient. Cells were washed twice prior to transfer. 10% of cells were retained for cellular analysis whereas the remaining 90% of cells were transferred into one non-irradiated *IgH*^*a*^ recipient mice by intravenous injection. A recall response was elicited by intravenously challenging mice with soluble NP-OVA 24 hours later.

### Flow cytometry

Single cell suspensions were prepared from bone marrow or spleen, erythrocytes lysed using an ammonium chloride-potassium solution, and lymphocytes isolated by using a Hisopaque-1119 (Sigma-Aldrich) density gradient. Cells were resuspended in PBS with 5% adult bovine serum and 2 mM EDTA prior to staining with antibodies and NP-APC. The following antibodies were purchased from Biolegend: 6D5 (CD19)-Alexa Fluor 700; GL7-FITC; 281-2 (CD138)-PE; RMM-1 (IgM)-APC; 11-26c.2a (IgD)-PerCP-Cy5.5 or -Brilliant Violet 605; 16-10A1 (CD80)-PE; RA3-6B2 (B220)-FITC, -Pacific Blue, or APC-Cy7; PO3 (CD86)-FITC; PK136 (NK-1.1)-PerCP-Cy5.5; M1/70 (CD11b)-Pacific Blue; HK1.4 (Ly-6C)-Brilliant Violet 510; 1A8 (Ly-6G)-Brilliant Violet 605; A7R34 (IL-7R)-Brilliant Violet 421; and E13-16.7 (Ly-6A/E)-PE. The following antibodes were purchased from eBioscience: 11/41 (IgM)-PerCP-e710; 11-26c (IgD)- FITC; 2B11 (CXCR4)-PerCP-e710; 2B8 (c-Kit)-PE-Cy7; and LG.7F9 (CD27)-APC. The following antibodies were purchased from BD Pharmingen: 53-6.7 (CD8a)-PE; RM4-5 (CD4)-PE-Cy7; A2F10.1 (CD135)-PE-CF594; and 93 (CD16/CD32)-PerCP-Cy5.5. Cells were stained on ice for 20 minutes. Germinal center B cells were enriched by staining cells with GL7-PE followed with anti-PE magnetic beads (0.5 uL/10^7^ cells, Miltenyi Biotec). Positive enrichment of GL7-expressing cells was performed using MACS LS columns (Miltenyi Biotec).

## Results

### ZBTB38 is highly expressed in B cell subsets

RNA-sequencing studies have reported expression of ZBTB38 in plasma cells [31]. To look more broadly across hematopoietic lineages, ZBTB38 gene expression in multiple cell types was first analyzed using available RNA-sequencing data assembled by the ImmGen Consortium [32]. A subset of cell types expressing DESeq2-normalized ZBTB38 counts greater than 800 are shown in **Figure 1A** [30]. High expression of ZBTB38 was observed in splenic plasma cells and plasmablasts (PC and PB), and in both light zone and dark zone germinal center B cells (LZ and DZ, **Figure 1A**). To confirm these data, wild-type mice were immunized with alhydrogel-adjuvanted 4- hydroxy-3-nitrophenyl-acetyl (NP) conjugated to ovalbumin (OVA) and naïve B cells (CD19^+^CD138^-^), NP-specific dark (CXCR4^+^) and light zone (CD86^+^) GC B cells (CD19^+^GL7^+^IgD^-^), and NP-specific splenic plasma cells (CD138^+^) were sorted 11 days after immunization [33]. RNA was extracted from sorted cells and quantitative RT-PCR performed to quantify ZBTB38 transcript levels. GC B cells and splenic plasma cells contained 9- and 3-fold, respectively, higher expression of ZBTB38 compared to naïve B cells (**Figure 1B, S1**). No differences in ZBTB38 expression were observed between light and dark zone GC B cells.

**Figure 1.**
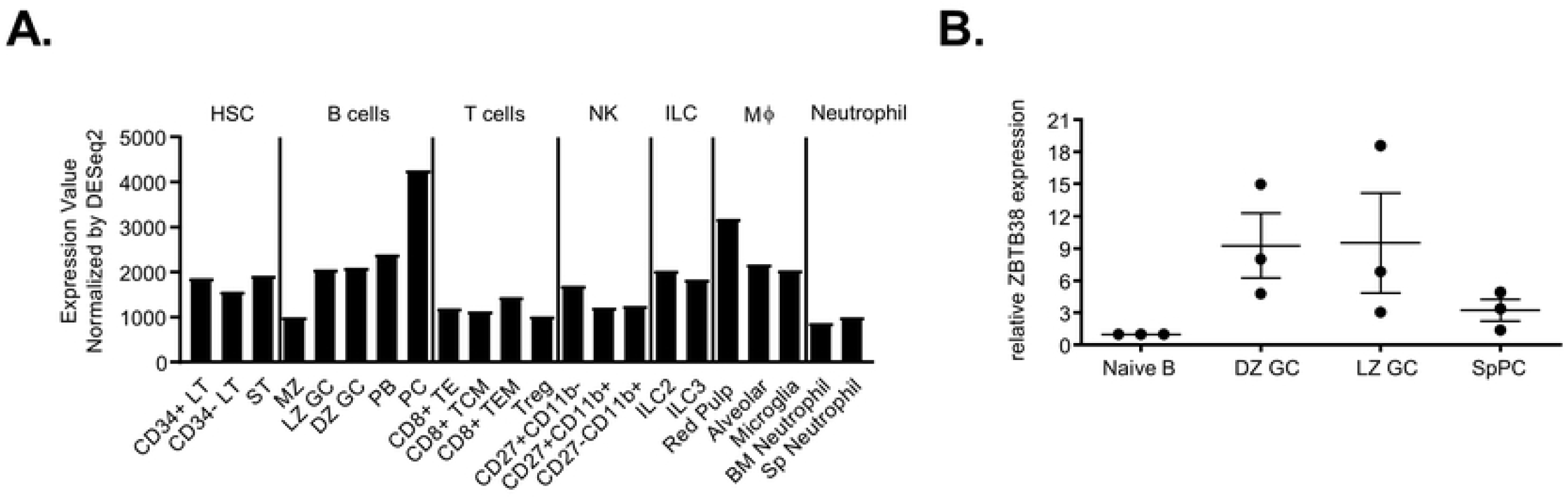
ZBTB38 is highly expressed by specific hematopoietic lineages. (**A**) ZBTB38 expression values in select cell subsets with DESeq2 count values over 800 were extracted from ImmGen’s RNA-seq SKYLINE and grouped based on cell type. (**B**) Naïve B cells, antigen-specific (NP^+^) dark (DZ, CXCR4^+^) and light (LZ, CD86^+^) zone germinal center B cells (GC, CD19^+^GL7^+^IgD^-^), and splenic plasma cells (SpPC, CD138^+^) were sorted 11 days after immunization of C57BL/6 mice with NP-OVA in alhydrogel, and Zbtb38 RNA levels quantified by quantitative RT- PCR. ZBTB38 expression was first normalized to 18S expression level followed by normalization to naïve B cells. Gating strategies are shown in S1 Fig. Mean ± SEM are shown; each symbol represents an individual mouse.

### Generation and validation of ZBTB38 conditional knockout mice

To assess the functional role of ZBTB38 *in vivo*, we generated *Zbtb38* floxed mice by targeting exon 3 (**Figure 2A**). This terminal exon contains the entire protein-coding sequence of ZBTB38. Single-stranded oligonucleotides containing loxP sites and flanking sequences of exon 3 of *Zbtb38* were microinjected alongside Cas9/guideRNA ribonucleoparticles into C57Bl6/J zygotes. gRNA sites were designed to flank the endogenous *Zbtb38* exon 3 and be disrupted upon homologous recombination with the targeting oligonucleotides. NdeI and EcoRI restriction sites were included in the oligonucleotides near the 5’ LoxP and 3’ LoxP sites to screen for successful homologous recombination of the targeting construct. Correct targeting of exon 3 was confirmed by PCR and restriction enzyme digestions (**Figure 2B**). Wild-type, targeted, and *Zbtb38*-deleted mice were distinguished using a set of three PCR primers flanking the 5’ and 3’ LoxP sequences (**Figure 2C**). To confirm deletion of *Zbtb38* exon 3 at the genomic level, *Zbtb38* f/f mice were crossed to mice expressing CMV-Cre to obtain germline ZBTB38 deletion. Tail genomic DNA from *Zbtb38* f/f x CMV-Cre was amplified and deletion was confirmed by a 4 kb reduction in PCR amplicon size (**Figure 2D**). Despite genome-wide association studies linking polymorphisms in the *Zbtb38* locus to human height [34-36], no differences were observed in the size of ZBTB38 deficient vs. wild-type littermates, and animals were born in expected Mendelian ratios (data not shown).

**Figure 2.**
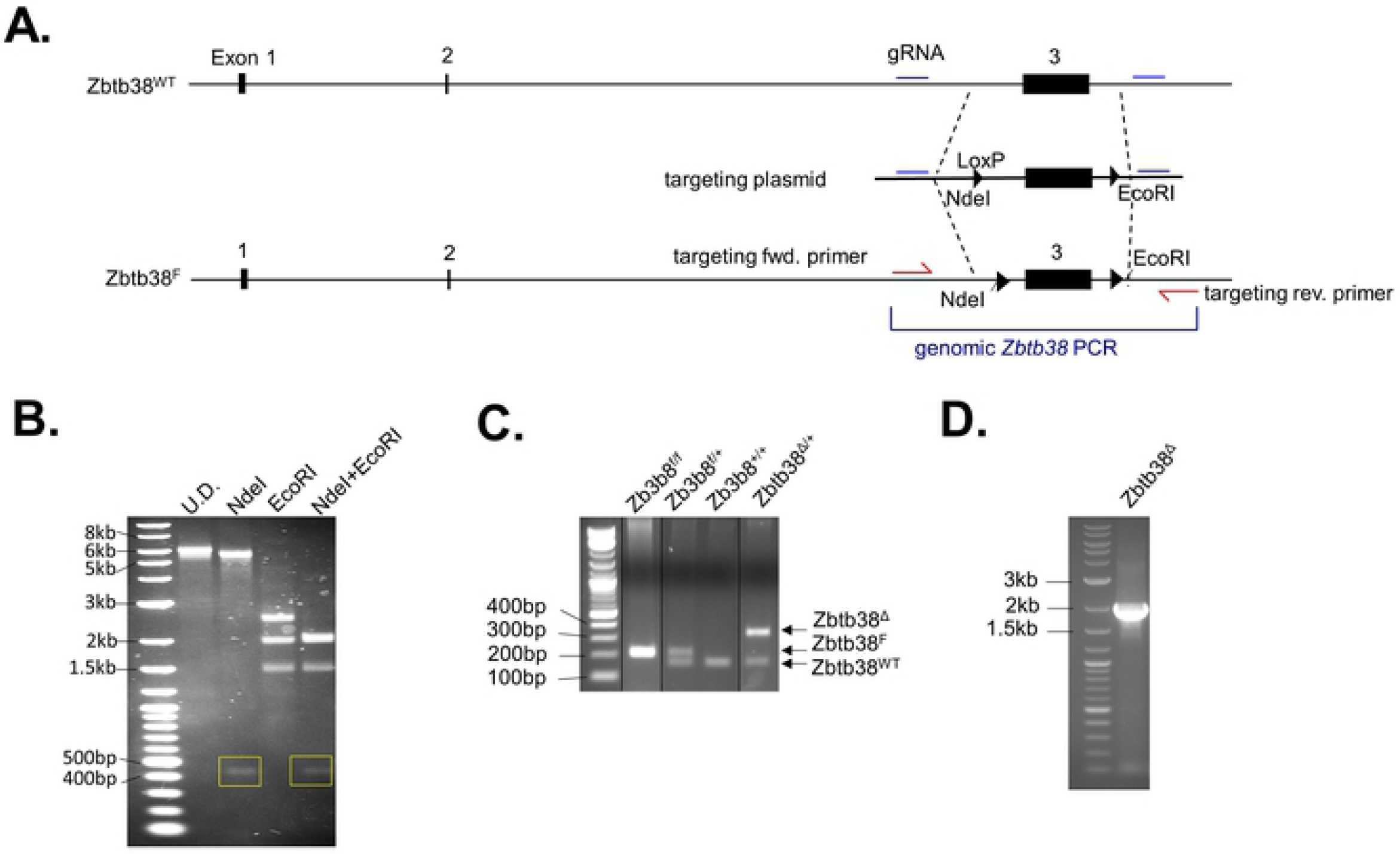
*Zbtb38* targeting strategy and confirmation of deletion. **(A)** Targeting strategy for exon 3 of *Zbtb38*. Guide RNA (gRNA) sites and targeting single-stranded oligonucleotides containing NdeI and EcoRI restriction sites were introduced to allow for screening of homologous recombination. **(B)** PCR strategy for exon 3 to confirm correct targeting. Lane 1 shows undigested PCR product, lane 2 shows NdeI digest (expected band sizes of 5.8 kb and 445 bp), lane 3 shows EcoRI digest (expected band sizes of 2.6 kb, 2 kb, and 1.6 kb), and lane 4 shows NdeI and EcoRI double digest (expected band sizes of 445 bp, 2.1 kb, 2 kb, and 1.6 kb). **(C)** Genotyping strategy and results to identify WT, floxed, or deleted *Zbtb38* alleles. **(D)** PCR confirming Zbtb38 deletion upon Cre expression. Primers used are identical to those used to amplify the genomic DNA as in **(B)**.

Given the high expression of ZBTB38 in blood lineages, we crossed *Zbtb38* f/f mice to VavCre mice (*Zbtb38* f/f x VavCre, ZBTB38 KO), which express Cre recombinase primarily in hematopoietic cells [37]. To confirm loss of ZBTB38 expression in this system, we performed RNA-seq on germinal center B cells isolated 2 weeks after immunization with NP-OVA. After mapping reads and aligning to the mouse genome, we observed that transcripts within the floxed exon 3 were completely abrogated in ZBTB38 KO mice, whereas intermediate levels were observed in *Zbtb38* f/+ x VavCre heterozygous littermates (**Figure 3A**) relative to *Zbtb38* f/f controls that lack Cre recombinase (ZBTB38 WT). We next used Salmon to quantify transcript abundances and DESeq2 to identify genes differentially expressed between ZBTB38- deficient mice and controls [29, 30]. Other than ZBTB38 itself, very few genes were dysregulated in ZBTB38-deficient cells (**Figure 3B**). To increase statistical power, we compared ZBTB38-deficient samples to both wild-type and heterozygous controls. Again, very few differences were observed (**Figure 3C**). Pathway analysis failed to reveal any transcriptional programs that were over- or under-represented in ZBTB38- deficient germinal center B cells (data not shown).

**Figure 3.**
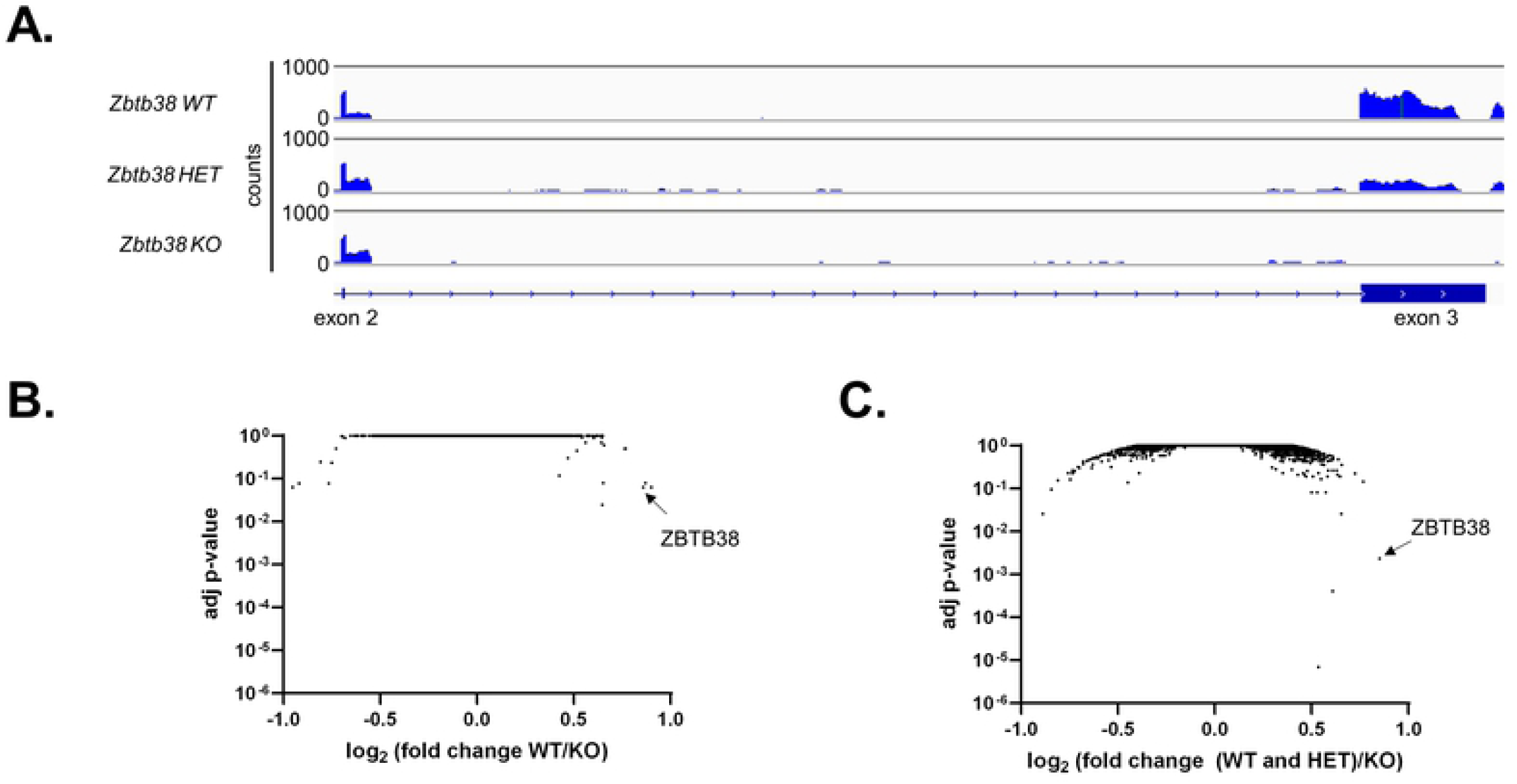
ZBTB38 deficiency minimally impacts gene expression in germinal center B cells. (**A**) RNA-seq reads across exons 2-3 of *Zbtb38*. Paired-end RNA-seq of antigen specific (NP^+^) germinal center B cells (CD19^+^GL7^+^IgD^-^) from *Zbtb38*^*fl/fl*^ (ZBTB38 WT), *Zbtb38*^*+/fl*^ *x Vav-Cre* (ZBTB38 HET), and *Zbtb38*^*fl/fl*^ *x Vav-Cre* (ZBTB38 KO) mice were aligned to the mm10 mouse genome and shown using IgV browser. (**B**) Volcano plot depicting differential gene expression between ZBTB38 WT (n=2) and ZBTB38 KO (n=6) mice. (**C**) Differential gene expression between ZBTB38 and ZBTB38 HET (n=4 in total) and ZBTB38 KO (n=6).

### ZBTB38 deficiency does not impair B cell responses

Deficiencies in two other BTB-ZF factors, ZBTB20 and ZBTB32, cause profound effects on plasma cell lifespan despite modest changes in gene expression [10-13]. Therefore, to determine if ZBTB38 has a functional role in primary B cell responses, we first examined GC reactions. We immunized ZBTB38 WT and KO mice with NP-OVA and quantified the frequency of NP-specific GC B cells 2 weeks later, which corresponds with peak GC reactions [38]. We observed no differences in the frequencies of NP-specific GC B cells between ZBTB38 WT and KO mice (**Figure 4A**). NP-specific serum antibodies were also similar between ZBTB38 WT and KO mice at all time points measured (**Figure 4B**). To specifically quantify the level of high affinity antibodies in the serum by ELISA, low density antigen (NP_4_) was used to probe for antibody binding. Low density NP (NP_4_) is used to capture antibodies with slow off-rates, which is correlated with antigen high affinity. This contrasts with high density NP (NP_16_), which can capture antibodies with faster off rates due to the increased concentration of antigen. Unlike the total levels of NP-specific antibodies over time, which plateaued two weeks after immunization, the concentration of high affinity antibodies increased steadily over time and plateaued four weeks after immunization (**Figure 4C**). No difference in the quantity of high affinity antibodies was observed between ZBTB38 WT and KO mice (**Figure 4C**). Furthermore, the extent of affinity maturation, quantified as the ratio of NP_4_ to NP_16_ endpoint titers, was similar between ZBTB38 WT and KO mice (**Figure 4D**).

**Figure 4.**
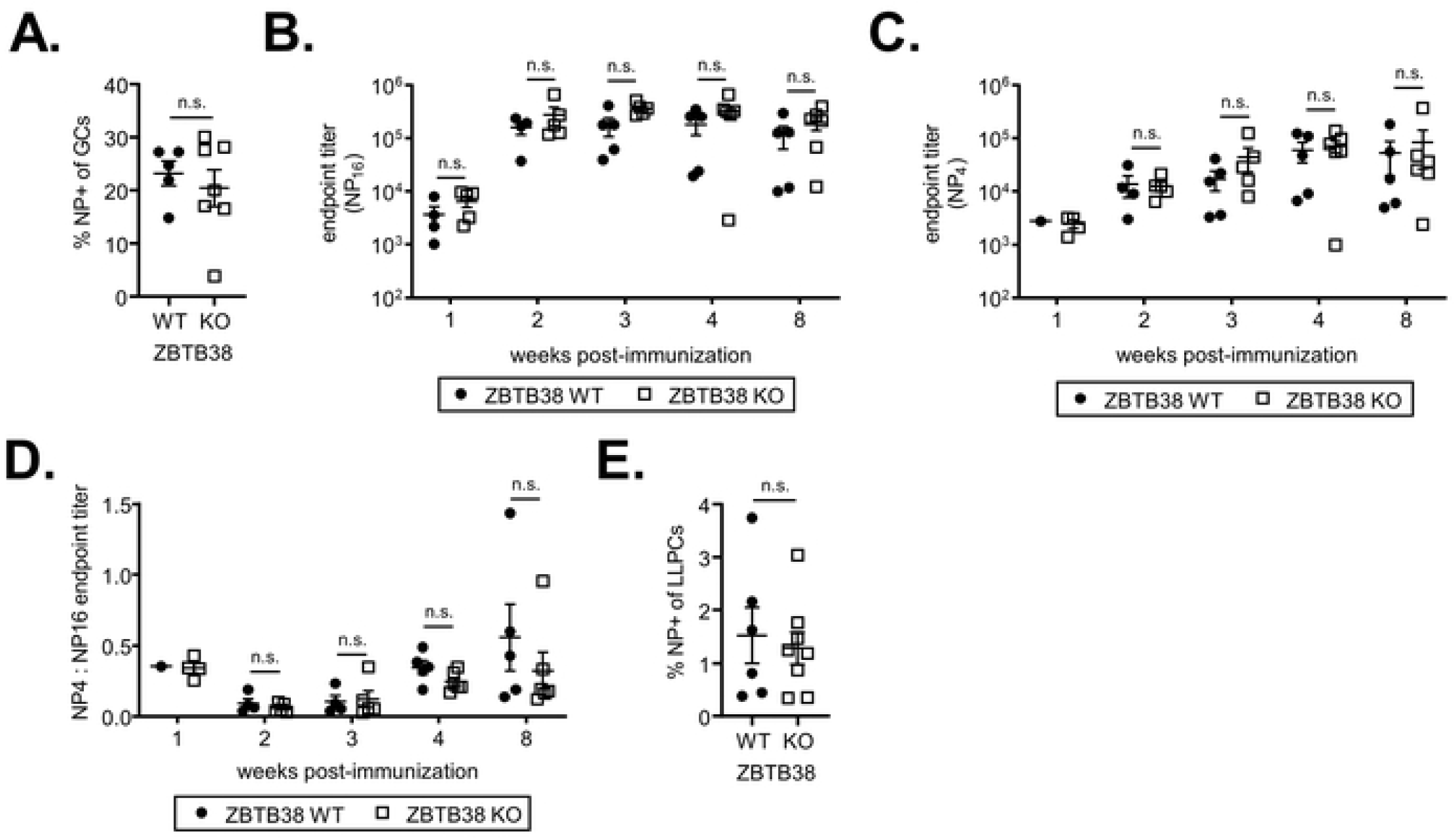
ZBTB38 is dispensable for primary B cell responses. (**A**) ZBTB38 WT and KO mice were immunized with NP-OVA and the frequency of NP-specific germinal center B cells two weeks post immunization was quantified by flow cytometry. Mean ± SEM are shown; each symbol represents an individual mouse. Statistical significance was calculated by unpaired student’s two-tailed t-test; n.s. = not significant (p > 0.05). (**B, C**) ZBTB38 WT and KO mice were immunized with NP-OVA and total serum antibody titers (**B**) and high affinity serum antibody titers (**C**) to NP were quantified by ELISA. Endpoint titers are calculated as the reciprocal serum dilution that was three standard deviations above background. Mean ± SEM are shown; each symbol represents an individual mouse. Statistical significance was calculated by Mann-Whitney test; n.s. = not significant (p > 0.05). (**D**) Affinity maturation of the antibodies was calculated as the ratio of endpoint titers to NP4: NP16 and plotted at each time point for ZBTB38 WT and ZBTB38 KO mice. Mean ± SEM are shown; each symbol represents an individual mouse. (**E**) The frequency of NP-specific long-lived plasma cells was calculated by flow cytometry. Long-lived plasma cells were analyzed 8 weeks post-NP immunization. Gating strategies are shown in S2 Fig. Mean ± SEM are shown; each symbol represents an individual mouse. Statistical significance was calculated by unpaired student’s two-tailed t-test; n.s. = not significant (p > 0.05).

Possible explanations for the similar serum antibody levels in ZBTB38 WT and KO mice include similar frequencies of antigen-specific LLPCs or compensatory increased antibody secretion from fewer LLPCs. To differentiate between these two possibilities, the frequency of NP-specific LLPCs was quantified by flow cytometry 8 weeks after alhydrogel-adjuvanted NP-OVA immunization of ZBTB38 WT and KO mice. The frequency of NP-specific LLPCs was similar between ZBTB38 WT and KO mice (**Figure 4E, S2**). These data demonstrate that ZBTB38 is dispensable for primary antibody responses to hapten-protein antigens.

To determine if ZBTB38 expression is required for secondary responses and MBC differentiation, the frequency of NP-specific memory B cells (CD19^+^GL7^-^IgM^-^IgD^-^ CD80^+^CCR6^+^) was quantified in ZBTB38 WT and KO mice 8 weeks after immunization with alhydrogel-adjuvanted NP-OVA. No difference in the frequency of NP-specific MBCs was observed (**Figure 5A, S3**). To assess MBC function, splenocytes from ZBTB38 WT and KO mice were adoptively transferred into allotype-distinct naïve IgH^a^ recipients and mice challenged with soluble NP-OVA 24 hours later. Donor IgH^b^ NP- specific antibodies originating from ZBTB38 WT and KO mice were tracked over time. NP-specific antibody titers were not altered by ZBTB38 deficiency (**Figure 5B**). Thus, ZBTB38 is also dispensable for secondary B cell responses.

**Figure 5.**
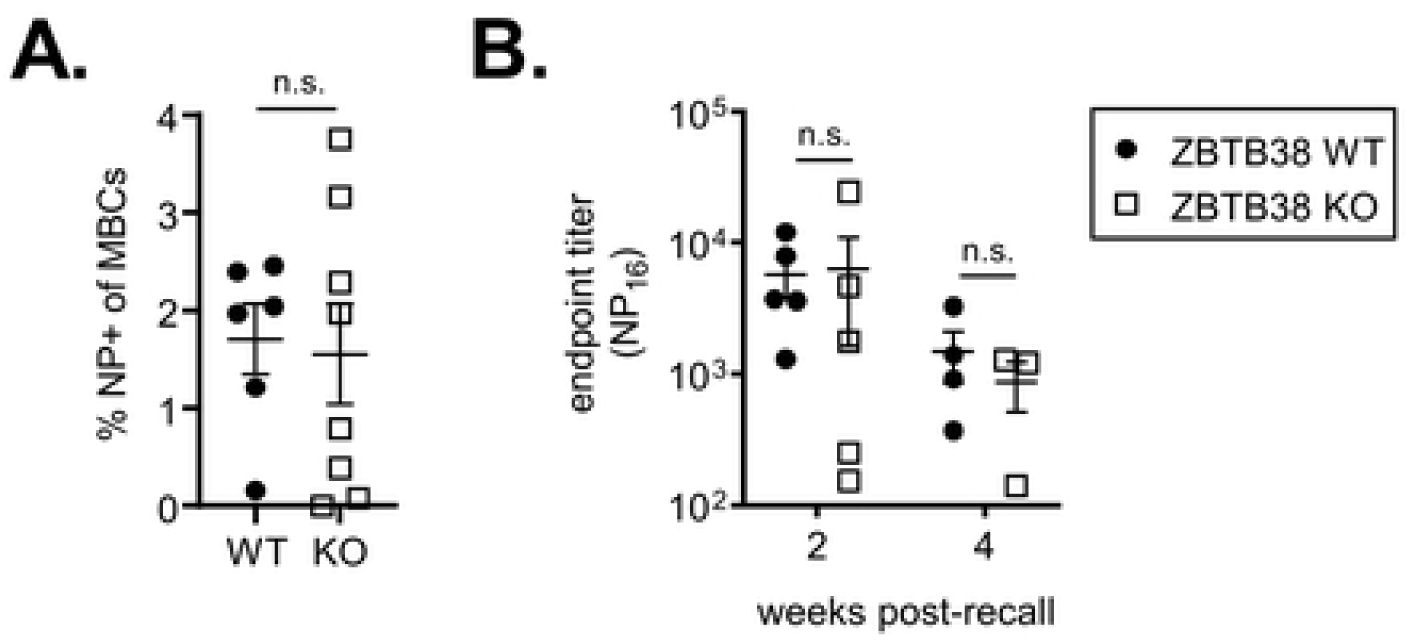
Memory B cell responses do not require ZBTB38. (**A**) The frequency of NP-specific memory B cells (CD19^+^GL7^-^IgM^-^IgD^-^CD80^+^CCR6^+^) was quantified by flow cytometry in ZBTB38 WT and KO mice 8 weeks after NP-OVA immunization. Gating strategies are shown in S3 Fig. Mean ± SEM are shown; each symbol represents an individual mouse. Statistical significance was calculated by unpaired student’s two-tailed t-test; n.s. = not significant (p > 0.05). (**B**) Splenocytes from ZBTB38 WT and ZBTB38 KO mice were adoptively transferred into IgHa naïve hosts and challenged with soluble NP-OVA one day after transfer. Donor (IgHb) antibody responses were calculated as the endpoint titer against high density NP. Mean ± SEM are shown; each symbol represents an individual mouse. Statistical significance was calculated by Mann-Whitney test; n.s. = not significant (p > 0.05).

### ZBTB38 deficiency does not alter the development of hematopoietic cells

Given ZBTB38 expression in hematopoietic progenitors (**Figure 1A**), we next assessed if ZBTB38 deficiency alters the development of lymphoid and/or myeloid lineages by quantifying the frequencies of different cell populations by flow cytometry. We first focused on hematopoietic stem cells (HSCs, cKit^+^Sca1^+^Flk2^-^CD27^+^) and progenitors with varying degrees of lineage commitment in the bone marrow [39]. HSCs differentiate into multipotent progenitors (MPPs, cKit^+^Sca1^+^Flk2^+^CD27^+^) that can give rise to both the myeloid and lymphoid lineages. MPPs can then differentiate into common myeloid progenitors (CMPs, cKit^+^Sca1^-^Flk2^+^FcγR^-^) or common lymphoid progenitors (CLPs, cKit^-/lo^Sca1^-^CD27^+^FcγR^-^Flk2^+^IL7Rα^+^) [40, 41]. CLPs give rise to B, T, natural killer (NK), and innate like cells (ILCs). CMPs give rise to basophils, eosinophils, mast cells, dendritic cells as well as granulocyte monocyte progenitors (GMPs, cKit^+^Sca1^-^Flk2^-^FcγR^+^) [41]. GMPs can then give rise to neutrophils and monocytes. We identified no statistically significant differences in the frequencies of these progenitor populations between ZBTB38 WT and KO mice (**Figure 6A, S4**).

**Figure 6.**
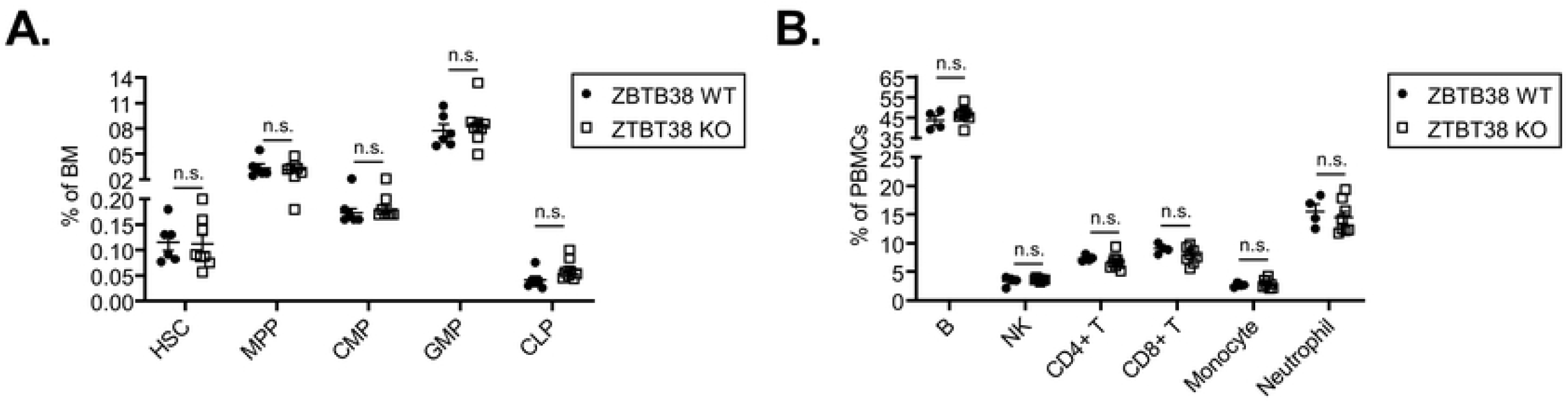
ZBTB38 deficiency does not impact hematopoietic development of maintenance. (**A**) Frequencies of hematopoietic stem cells (HSCs, cKit^+^Sca1^+^Flk2^-^ CD27^+^), multi-potent progenitors (MPPs, cKit^+^Sca1^+^Flk2^+^CD27^+^), common myeloid progenitors (CMPs, cKit^+^Sca1^-^Flk2^+^FcγR^-^), granulocyte monocyte progenitors (GMPs, cKit^+^Sca1^-^Flk2^-^FcγR^+^), and common lymphoid progenitors (CLPs, cKit^-/lo^Sca1^-^ CD27^+^FcγR^-^Flk2^+^IL7Rα^+^) in the bone marrow from ZBTB38 WT and KO mice were quantified by flow cytometry. Gating strategies are shown in S4 Fig. Mean ± SEM are shown; each symbol represents an individual mouse. (B) Frequencies of B cells (B220^+^), natural killer (NK, NK1.1^+^) cells, CD4^+^ and CD8^+^ T cells (B220^-^CD11b^-^NK.1^-^), monocytes (CD11b^+^Ly6C^hi^Ly6G^-^), and neutrophils (CD11b^+^Ly6C^+^Ly6G^+^) of peripheral blood mononuclear cells (PBMCs) from ZBTB38 WT and KO mice were quantified by flow cytometry. Mean ± SEM are shown; each symbol represents an individual mouse. Statistical significance was determined by unpaired student’s 2-tailed t-test; n.s. indicates no significance (p > 0.05).

To assess if mature hematopoietic lineages require ZBTB38 for their development or maintenance, we analyzed the frequencies of different peripheral blood mononuclear cell (PBMCs) populations. We observed no differences between ZBTB38 WT and KO mice in the frequencies of B cells (B220^+^), NK cells (NK1.1^+^), T (CD4^+^ or CD8^+^ B220^-^NK1.1^-^CD11b^-^), monocytes (CD11b^+^Ly6C^hi^Ly6G^-^), or neutrophils (CD11b^+^Ly6C^+^Ly6G^+^) (**Figure 6B**). Thus, ZBTB38 is not required for the development or maintenance of these hematopoietic lineages.

## Discussion

BTB-ZF family members such as BCL-6, ZBTB32, and ZBTB20, have critical roles in different aspects of B cell responses. ZBTB38, another member of the BTB-ZF family, has been implicated in DNA damage responses and replication efficiency. This occurs through repression of MCM10 expression, and potentially binding of other methylated CpG sites throughout the genome [18, 20]. Replication fidelity and DNA damage responses are key processes during germinal center reactions, as B cells accumulate somatic mutations and undergo dsDNA breaks as part of immunoglobulin isotype switching [42]. Yet our data demonstrate that ZBTB38 is dispensable for both primary and recall B cell responses to model T-dependent antigens despite high levels of expression in both germinal center B cells and antibody-secreting plasma cells. ZBTB38 is also dispensable for maintaining homeostasis of lymphoid and myeloid hematopoietic lineages. Thus, the evolutionary and functional reasons why ZBTB38 is expressed in the hematopoietic and immune system are not fully resolved. Instead, the most important roles for ZBTB38 may lie in other cell types, tissues, and/or physiological contexts.

Genome-wide association studies in humans have identified single nucleotide polymorphisms (SNPs) in ZBTB38 that are associated with shorter stature in Chinese populations but taller stature in Korean populations [34-36, 43]. However, when *Zbtb38* f/f mice were crossed to CMV-Cre expressing mice, germline deletion of ZBTB38 did not result in observable differences in the length or weight of the mice (data not shown). Further characterization of how SNPs influence ZBTB38 function, and identifying the location of SNPs, may provide further explanation of why certain polymorphisms are associated with altered height.

ZBTB38 deletion or knockdown has resulted in impaired cellular processes in neuronal injuries or tumors. For instance, increasing ZBTB38 expression reduces apoptosis and promotes autophagy in a spinal cord injury model, and results in increased neuronal repair [25, 44, 45]. In contrast, ZBTB38 expression has been shown to promote proliferation and differentiation of a neuroblastoma cell line [46]. These differences in the functional roles of ZBTB38 may be attributed to subtype-specific sensitivity of neurons to oxidative stress [47]. ZBTB38, along with USP9X, a deubiquitinase, is required to limit basal reactive oxidative species (ROS) levels and the response to oxidative stress [48]. Given the high levels of ZBTB38 expression in neurons, perhaps ZBTB38 is involved with balancing neuronal death with recovery after various challenges. Neuronal insult, such as injury, stroke, or cancer may be necessary to identify processes regulated by ZBTB38. Future studies focused on such other cell types and systems will be facilitated by the novel *Zbtb38* f/f mice that we have generated.

## Acknowledgments

This work was supported by National Institutes of Health grant R01AI099108 (D.B.). R.W. was supported by the National Science Foundation Graduate Research Fellowship Program (DGE-1143954). The authors thank M. White in the Transgenic and Knockout Mouse Core at Washington University for assistance with microinjections. Flow cytometry experiments reported in this publication were supported by Flow Cytometry Shared Resource at the University of Arizona Cancer Center and the National Cancer Institute of the National Institutes of Health under award number P30CA023074.

## Author Contributions

Conceived and designed the experiments: RW DB. Performed the experiments: RW. Analyzed the data: RW DB. Wrote the paper: RW DB.

## Supporting Information

**S1 Figure. Gating strategy for splenic plasma cells, light zone germinal center B cells, and dark zone germinal center B cells**. Flow cytometric gating strategies for NP-specific splenic plasma cell (SpPC), light zone (LZ) and dark zone (DZ) germinal center (GC) B cells shown in Figure 1B.

**S2 Figure. Gating strategy for long-lived plasma cells**. Flow cytometric gating strategy for NP-specific long-lived plasma cells (LLPCs) in the bone marrow.

**S3 Figure. Gating strategy for isotype-switched memory B cells**. Flow cytometric gating strategy for NP-specific, isotype-switched memory B cells (swIg MBCs) in the spleen. Cells were gated on CD19^+^GL7^-^.

**S4 Figure. Gating strategy for bone marrow progenitors**. Flow cytometric gating strategies for bone marrow progenitors shown in Figure 6A. HSC, hematopoietic stem cell; MPP, multi-potent progenitor; CMP, common myeloid progenitor; GMP, granulocyte monocyte progenitor; CLP, common lymphoid progenitor.

